# To catch a hijacker: abundance, evolution and genetic diversity of P4-like bacteriophage satellites

**DOI:** 10.1101/2021.03.30.437493

**Authors:** Jorge A. Moura de Sousa, Eduardo P.C. Rocha

## Abstract

Bacteriophages (phages) are bacterial parasites that can themselves be parasitized by phage satellites. The molecular mechanisms used by satellites to hijack phages are sometimes understood in great detail, but the origins, abundance, distribution, and composition of these elements are poorly known. Here, we show that P4-like elements are present in more than 10% of the genomes of Enterobacterales, and in almost half of those of *Escherichia coli*, sometimes in multiple distinct copies. We identified over 1000 P4-like elements with very conserved genetic organization of the core genome and a few hotspots with highly variable genes. These elements are never found in plasmids and have very little homology to known phages, suggesting an independent evolutionary origin. Instead, they are scattered across chromosomes, possibly because their integrases are often exchanged with other elements. The rooted phylogenies of hijacking functions are correlated and suggest longstanding co-evolution. They also reveal broad host ranges in P4-like elements, since almost identical elements can be found in distinct bacterial genuses. Our results show that P4-like phage satellites constitute a very distinct, widespread and ancient family of mobile genetic elements. They pave the way for studying the molecular evolution of antagonistic interactions between phages and their satellites.

## Introduction

Bacteriophages (phages) use their bacterial hosts to replicate, which usually ends in bacterial death and the release of virions containing the phage genome. However, phages have their own parasites, satellite mobile elements that cannot produce virions and instead hijack those of functional (so-called helper) phages [1]. The best studied phage satellites are P4 in *Escherichia coli* [2,3], *Staphylococcus aureus* pathogenicity islands and phage-inducible chromosomal islands (SAPIs and PICIs) in *Bacillales* and enterobacteria [4], and phage-inducible chromosomal islands-like elements (PLEs) in *Vibrio* spp [5,6]. Their genome sizes vary between 11 kb in P4 and 18 kb in PLE. These values are well below the average genome size of dsDNA temperate phages, which in these clades tend to be larger than 30 kb [7]. Phage satellites integrate the bacterial host genome and may change its phenotype. For example, some SaPIs encode important virulence factors [8], whereas P4 and PLE encode anti-phage defence systems [9,10].

The satellite-helper system that is best understood in molecular terms is the pair P4-P2. Decades of studies have revealed the key functions of P4 genes [3,11]. P4 was initially regarded as a phage-plasmid, i.e. a phage that can reside in the lysogenic cells as a plasmid [11], and to-date there is still no clear evidence on whether P4 is phage, a plasmid or a completely distinct mobile element. While its *cnr* gene controls the copy-number of the plasmid stage, P4 also encodes an integrase and is often found integrated in the chromosome. Its transmission through the plasmid lifecycle is thought to be infrequent, as only ~1% of the P4 infections result in its establishment as a plasmid [12]. The co-infection of an *E. coli* bacterium with both P2 and P4 provides the context for the parasitism of the latter. P4 subverts the gene expression of P2 using Ash (also called ɛ), which inactivates the repressor of the helper phage, causing its induction [13]. The *δ* gene is an homolog of P2’s *ogr* transcriptional activator and both promote the expression of P2’s and P4’s late genes [14]. The *α* gene encodes a protein with primase and helicase activities and is essential for DNA replication [15]. The gene *alpA* is predicted to act as a DNA-binding transcriptional regulator. Finally, the concerted action of the Psu and Sid proteins results in the constriction of P2 capsids and allows P4 to hijack the P2 virion [16,17]. As a result, the virions are able to package the DNA from the small genome of P4, but not from the larger genome of P2. Recent structural work has revealed that Sid and Psu are homologs with some sequence similarity and similar folds [18].

Despite the exquisite knowledge obtained in the past decades on the molecular interactions between P4 and P2, we know little about the diversity and distribution of the P4 family of satellites. P4-like elements may be difficult to identify using prophage identification tools because they lack typical distinguishing phage components, like packaging or virion proteins, and the diversity of their gene repertoires is unknown. Nevertheless, a few studies have suggested the existence of other P4-like elements. For instance, a cryptic P4-like prophage was found to excise and express an AlpA homolog [19], and another one, encoding a retron, was shown to require P2 for transfer [20]. Additionally, a study of prophage domestication identified 28 elements with a homolog of *sid* [21], and a recent study identified more than 5000 homologs of *psu* in ca. 20000 *E. coli* genomes [9].

These studies raised key questions. Is P4 part of a family of mobile genetic elements with conserved gene repertoires and genetic organizations? What is the core genome and the genetic organization of such a family? Are members of the family abundant in bacterial genomes? Have P4-like elements recently derived from other mobile genetic elements, e.g., phages, or are they ancient? To provide answers to these questions, we searched bacterial genomes for regions with clusters of homologs to the abovementioned key components of P4. Given the lack of available methods to detect P4-like satellites, we studied the composition of these clusters to uncover and characterize a putative family of P4-like satellites. We quantified their abundance, which resulted in the identification of ca. 1000 novel P4-like mobile elements. This allowed us to study their prevalence and association with bacterial hosts. We also characterized the composition and organization of the gene repertoires of P4-like elements, as well as the phylogenetic history of key genes. Our results uncover the hidden diversity of P4-like satellites and highlight their role as a distinct mobile element in the microbial world.

## Results

### P4-like satellites are a large and well-defined family of mobile elements

The genetic diversity of P4-like satellites is currently unknown, which complicates their identification. We started our study by searching for genomes related to P4 amongst the 2487 sequenced phages from the NCBI RefSeq database, using the weighted Gene Repertoire Relatedness index (wGRR, see Methods), a measure of the proteome similarity between pairs of genomes. This failed to reveal P4-like elements, since 93% of the phages are completely unrelated to P4 (wGRR=0, Fig S1A), whilst the remainder 7% show very low wGRR (<0.06). The analysis of sequence similarity between P4 proteins and all the proteins in the phage database revealed hits with an e-value higher than 10^-5^ for the integrase, the replicase and the transcriptional regulators. Delta proteins showed low homology (all hits with less than 40% identity) with a few P2-like genomes. Moreover, amongst the 26984 prophages detected in 13513 bacterial genomes, less than 0.1% have a wGRR with P4 above 0.1 (Fig S1B). The few prophages with high wGRR consisted of a P4-like element close to a large prophage. We conclude that there are no phages and very few prophages with extended homology to P4. Hence, either P4-like elements are extremely rare or they are missed by current computational approaches for phage identification.

We used publicly available and custom-built Hidden Markov Models (HMM) profiles to search for homologs of P4 key components – integrase, *psu, δ*, *sid*, *alpA*, *ε* and *α* - in bacterial genomes, including 22310 plasmids (see Methods). We ordered these homologs by their genomic location and clustered them in *sets*: groups of co-localized homologs to these components (i.e., less than 10 genes apart, Fig S2). Since we were unaware if these components would be present in all elements, we allowed sets to miss one or two components. We found 1037 sets (Table S1), among which 502 contain all the seven components of P4 (hereafter referred to as Type A, Fig 1A), 456 lack one component (Type B) and only 79 lack two (Type C). Each possible combination of missing components was deemed as a variant of each type (Fig 1A). Among the Type B sets, most (56%) lack an homolog to *alpA* (TypeB#03). Variants lacking *α*(TypeB#02) or *psu* are also frequent (TypeB#06). This suggests that these components are either facultative or can be replaced by functional homologs in some P4-like elements. On the other hand, only one variant missing *δ*(TypeB#05) was found, suggesting that this is a conserved and essential component of functional P4-like satellites. The most abundant Type C variant lacks both *ε* and *α*(TypeC#01). Noticeably, seven of the 21 possible variants of Type C were never found. The abrupt decline in the abundance of Type C elements, relative to Types A and B, suggests that there is a clear distinction between elements with a common set of key components homologous to P4 and the other mobile genetic elements.

**Fig. 1.**
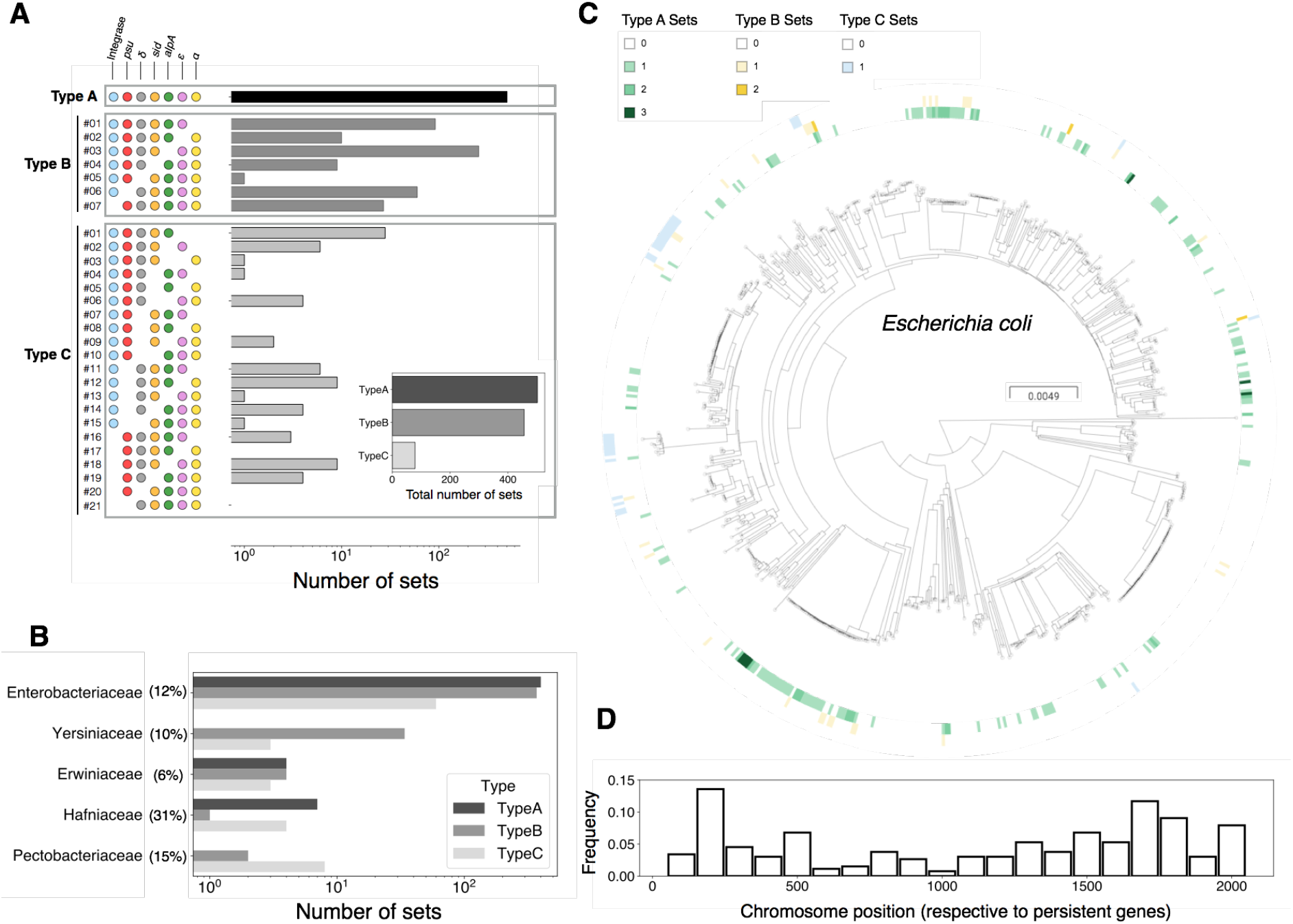
The abundance, genetic organization and bacterial hosts of P4-like elements. **A)** Log-scale abundance of the variants, defined by the combinations of core components (circles to the left of the distributions). The inset shows the total abundance per type, considering all variants. **B)** Distribution of the different sets in bacterial families. The percentages following the families indicate the fraction of bacterial genomes of that family that encode P4-like satellites. **C)** Distribution of P4-like sets in the *E. coli* species tree. The tree was built using Maximum Likelihood, using the alignment of the core genome (2107 genes) from 657 *E. coli* genomes obtained using PanACoTA [23] and FastTree with default parameters [24]. The presence and abundance of sets of Type A (green), B (yellow) or C (blue) are shown in the inner, middle and outer circle, respectively, with genomes that encode for more than one set of either type shown as a stronger shade of their respective color. **D**) Position of the P4-like element along the genome of *E. coli.* The core genes of the species were ordered and numbered in function of the position of the gene in the chromosome of strain MG1655 from 1 to 2107. Positions between core genes define genome intervals. The position indicates the interval where the P4-like element was identified. Hence, an element integrating the same region across genomes would have the exact same position.

There has been a longstanding uncertainty concerning the relevance of the plasmid stage in P4 [11,22]. This prompted us to analyze whether the sets we identified – Types A, B or C – are located in bacterial chromosomes or in plasmids. All sets were found in bacterial chromosomes. Although we found some isolated components, or rare sets of two components (one being often an integrase) in extrachromosomal replicons, we could not find a plasmid with more than three components. This suggests that P4-like satellites are not plasmids under normal physiological conditions.

We then analysed the taxonomic distribution of the bacteria with P4-like elements (Types A, B and C). The genomes of Enterobacteriaceae include 92% of the elements, and these are found in 12% of their genomes (Fig 1B and Fig S3). Most genomes have one element, but some have up to three (Fig 1C and Fig S4). The most represented species in our genome dataset is *E. coli*. In this species, 44% of the genomes (out of 657) encode at least one P4-like element. These elements are dispersed and prevalent across the species tree (Fig 1C). To confirm that their prevalence is not the result of a single ancestral infection, we analysed the positions of P4-like elements in the chromosomes of the species (see Methods). We located the elements in relation to the positions of the neighboring core genes and found that P4-like elements are scattered across the *E. coli* chromosome (Fig 1D). The dispersion of the elements across the phylogenetic tree and across the chromosome shows that these elements have proliferated by horizontal transfer across the species.

We also found P4-like elements in genomes of bacterial families that are poorly represented in the genome database: Yersiniaceae (4% of the elements), Hafniaceae (1.3%), Erwiniaceae (1.2%) and Pectobacteriaceae (1%). Interestingly, the P4-like elements in the Yersiniacea family are exclusively comprised of Type B that lack a *psu* homolog (TypeB#06). Even though estimates of the prevalence of P4-like elements in these poorly sampled bacterial families are less reliable, given the few genomes available for some families, our analysis reveals that between 6% and 31% of the genomes of each family encode at least one element (Fig 1B). In summary, P4-like satellites are a very abundant and characteristic mobile element, widespread in the genomes of Enterobacterales.

### Genomic characterization of the P4-like satellite family

To study the genetic organization of P4-like elements, we analyzed the order of the components using the integrase as a starting position. These analyses were restricted to Types A and B, which comprise ~92% of the P4-like satellites and are more likely to include functional elements. The order of the components is very similar for Types A and B, and 77% of the former have the organization of P4 (Fig 2A and Fig S5-6). The most frequent exception concerns a swap in the order between *psu* and *δ*(20% of the total), whilst in a few others the homologs of *alpA* follow *α*’s homologs (1.2%). In very rare cases (<1%), the integrase appears next to *α*, and the order of the rest of the components is reversed relative to P4. Such cases seem to reflect the presence of other mobile genetic elements encoding their own integrase and neighboring P4-like elements whose integrase is either missing or more than 10 genes apart from *psu*. The distance between consecutive components is diverse (Fig 2A, boxen plots). The genes of some of the core components, like the operon formed by *psu*, *δ* and *sid,* tend to be adjacent, whilst the number of genes between the integrase and *psu* ranges from one to nine, if we consider only the variants where the integrase precedes *psu* (in all cases the outliers in the distributions correspond to the rare organizational variants). Overall, these results suggest that the organization of the P4-like genome is very conserved.

**Fig. 2.**
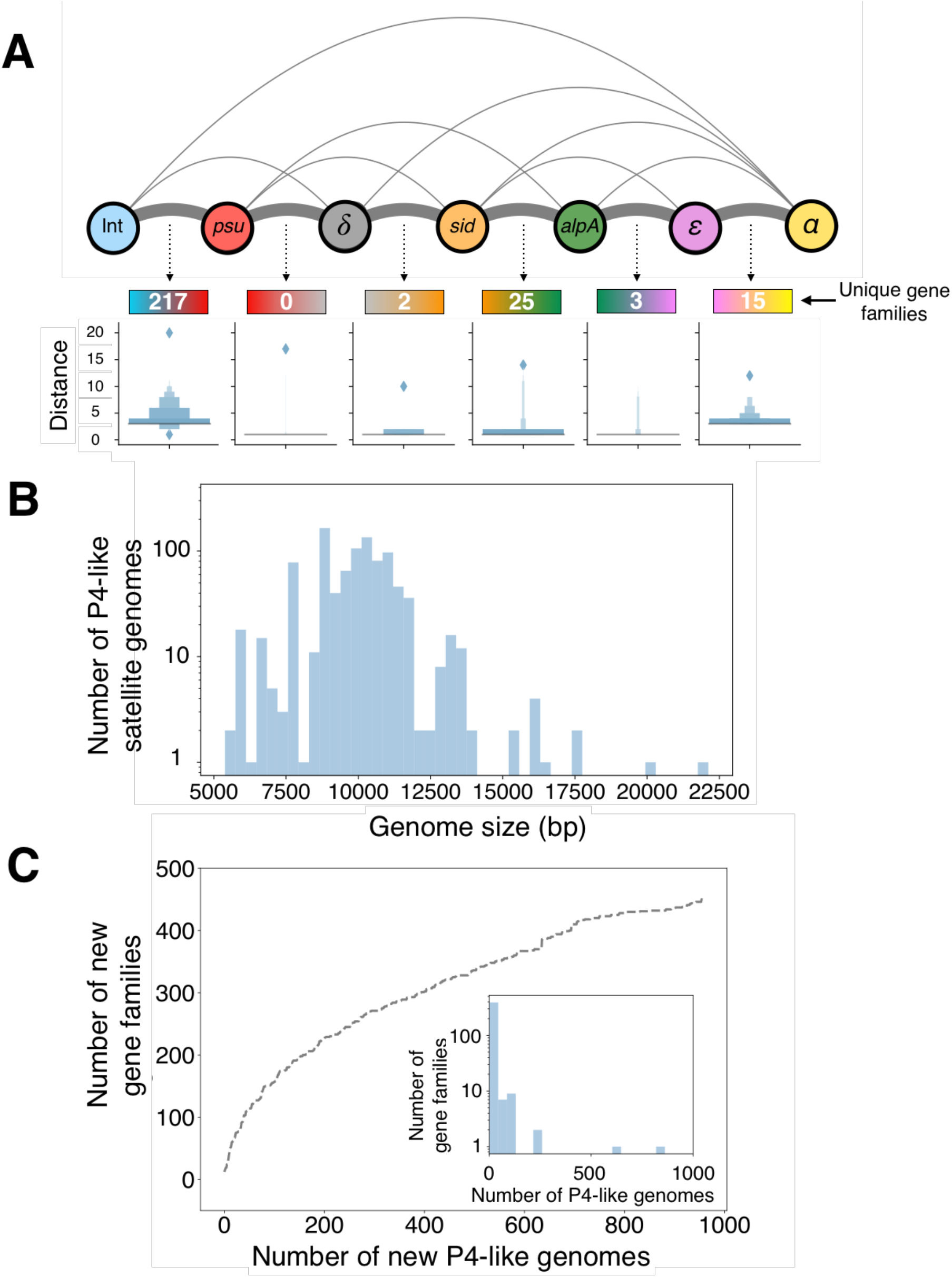
The genomes and pangenome of P4-like satellites. **A)** Genetic organization of the P4-like elements. The thickness of the edges is proportional to the frequency of adjacency of pairs of components. Below the major edges is indicated the number of gene families between the two genes (for Type A elements with the most common organizational variants only). The box plots show the distribution of the number of open reading frames between consecutive components. The outliers of these distributions represent the rare organizational variants where the most frequent pairs of components are further than 10 genes apart. **B)**Histogram of the genome size of P4-like elements. **C)**Rarefaction curve of the gene families of P4-like genomes, excluding those of core components. New gene families were sequentially added from randomly ordered P4-like genomes. The inset shows the frequency of the gene families.

Since the core components are nearly ubiquitous and organized similarly, we can use the integrase and *α* homologs to tentatively delimit P4-like elements. In P4, these two genes are separated only by the *att* site, with no other genes between them. Further, we could not find prevalent gene families (i.e., present in more than 25% of the elements) in the five genes before the integrase or the five genes after *α*, across all sets. For the few Type B sets lacking *α* and for the rare organizational variants where *α* is not the last component in the set, we used the last component to delimit the satellite genome. While we cannot exclude that some elements may be larger, our results suggest the two components are suitable to delimit the P4-like satellites.

The genomes of P4-like satellites have a median of 10 kb (Fig 2B), close to the 11 kb of P4, encoding ca. 11 proteins (Fig S7). However, there is considerable variation in their sizes. Some elements have close to 5 kb, whereas a very small number (11 genomes) are more than 15 kb long. The smaller genomes mostly correspond to Type B variants lacking either the integrase or *α*, and were delimited by *psu* and *ε*, which may justify their small size. The removal of these cases raises the minimal genome to ~7 kb (Fig S8), but still leaves considerable variation in size. The largest elements are often tandem mobile elements or groups of genes for P4-like components adjacent to complete P4-like elements. They may reflect pseudogenization of tandem elements. The conservation of the repertoires of core components and the large variation in size of the elements suggests the existence of high genetic diversity within non-core components. We computed the pangenome of the P4-like family to assess this point more precisely. After removing the core components mentioned above, the gene family frequency spectrum is L-shaped with most gene families (381, 85%) present in fewer than 10 genomes. This abundance of low frequency gene families results in an open pangenome (Fig 2C), suggesting that P4-like elements have a large gene repertoire. The estimation of its full size and diversity will require sequencing further genomes in a more varied range of species. We then computed the gene families specific to the locations between each pair of core components. We restricted this analysis to genomes of Type A with the most conserved genetic organization, in order to characterize the local pangenome using only the common pairings of adjacent components. Most gene families observed between consecutive core components are located between the integrase and *psu* or between *sid* and *alpA* (Fig 2A, numbers below organization diagram). The first region corresponds to the previously described hotspot of phage defense systems [9], and includes the *cos* site in P4. The region between *ε* and *α* has two frequent gene families. One is present in 90% of the elements and includes the *cnr* gene of P4. The other is present in 61% of the elements and encodes a small protein (75 aa) of unknown function that is absent from P4. This suggests that P4 lacks a protein that is prevalent in its family of satellites.

To compare the gene repertoires of P4-like elements, we computed the wGRR between them. Most comparisons (81%) revealed wGRR values between 0.6 and 0.2, suggesting that most P4-like genomes have considerable genetic differences in spite of the core genes present in most elements (Fig 3A). The peak at wGRR close to one represents pairs of elements that are practically identical and may result from vertical descent or recent transfer across bacteria. We used hierarchical clustering on the wGRR pairwise comparisons matrix to identify groups of similar satellites. The large groups contain both Type A and B elements, but some sub-clusters set them apart, suggesting the existence of relatively recent lineages with different repertoires of core components. The largest clusters are those of satellites detected in *Escherichia* (in two different clusters), *Klebsiella* and *Salmonella*. P4-like satellites detected in the *Yersiniaceae* family (all from Type B) also form a cluster. This suggests that elements tend to cluster in function of their bacterial host species or genus. Nevertheless, 4% of the pairwise comparisons with high wGRR (≥0.9, N=19961) concern elements present in different bacterial species, often in different genus. Furthermore, 14% of the 5731 pairs of elements with average nucleotide identity (ANI) higher than 95% (the threshold typically used to separate bacterial species [25]) are found in different bacterial species. This suggests that P4-like satellites can disseminate across distantly related bacteria.

**Fig. 3.**
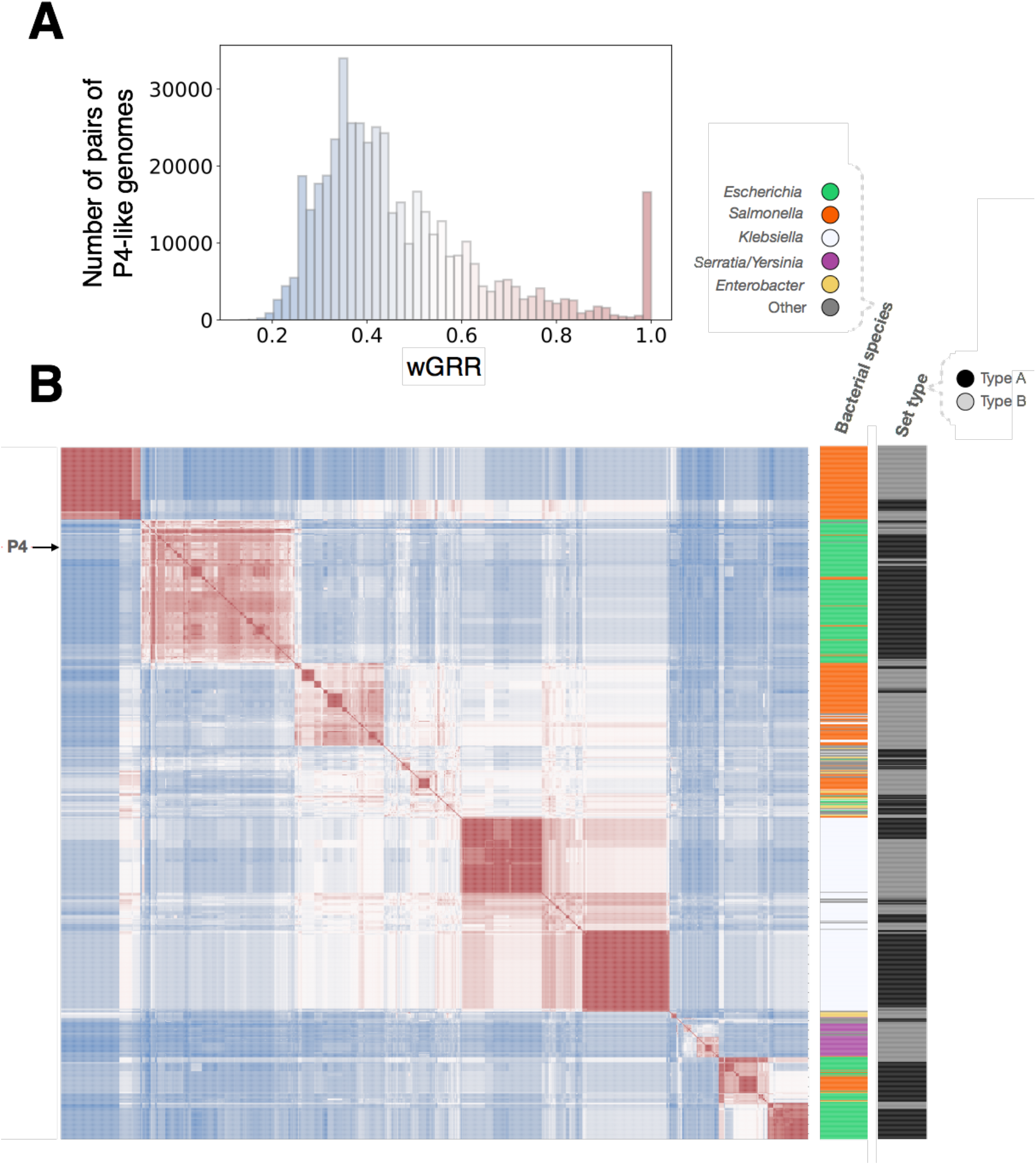
Comparison of gene repertoires of P4-like satellites. **A)** Histogram of wGRR for all pairwise comparisons between the 958 P4-like genomes (of Types A and B). **B**) Heatmap of the matrix of the wGRR values ordered using hierarchical clustering. The colors follow the same code as in **A**. The columns to the right of the heatmap indicate the bacterial species where the P4-like genome was detected and the type (A or B) of the P4-like genome. The position of the P4 genome in the matrix is indicated on the left side.

### Evolution of P4-like satellites

The genes *psu* and *sid* encode key functions for the hijacking of P2 virions by P4. These genes are structural homologs and share some sequence similarity [18], which suggests they derived from the same ancestral protein. In such cases, the joint phylogeny of two proteins allows to root the sub-tree of each of them [26]. To investigate the evolutionary history of *psu* and *sid*, we aligned their proteins from sets of Type A, B and C and used it to infer the phylogeny. The resulting tree (File S3) is well supported at most key nodes and shows two distinct monophyletic clades that correspond to either Psu or Sid (Fig 4). Hence, it can be used to understand the evolution of the two proteins and to distinguish ancestral from derived states. We have also built phylogenetic trees for *δ* and the integrase, and computed the patristic distances (*i.e.*, the sum of the lengths of branches that link two nodes in a tree) between the proteins of each family. The correlation between patristic distances of the four protein families show diverse patterns (Fig S9, Files S4-S7). Notably, the Spearman correlations between the patristic distances of the integrase and those of the other components are very low (between 0.01 and 0.16), showing that this protein has a very different evolutionary history among P4-like elements. The correlations between *δ*, Psu and Sid are much higher, especially those between the two latter proteins (R = 0.76), revealing joint evolution within the elements.

**Fig. 4.**
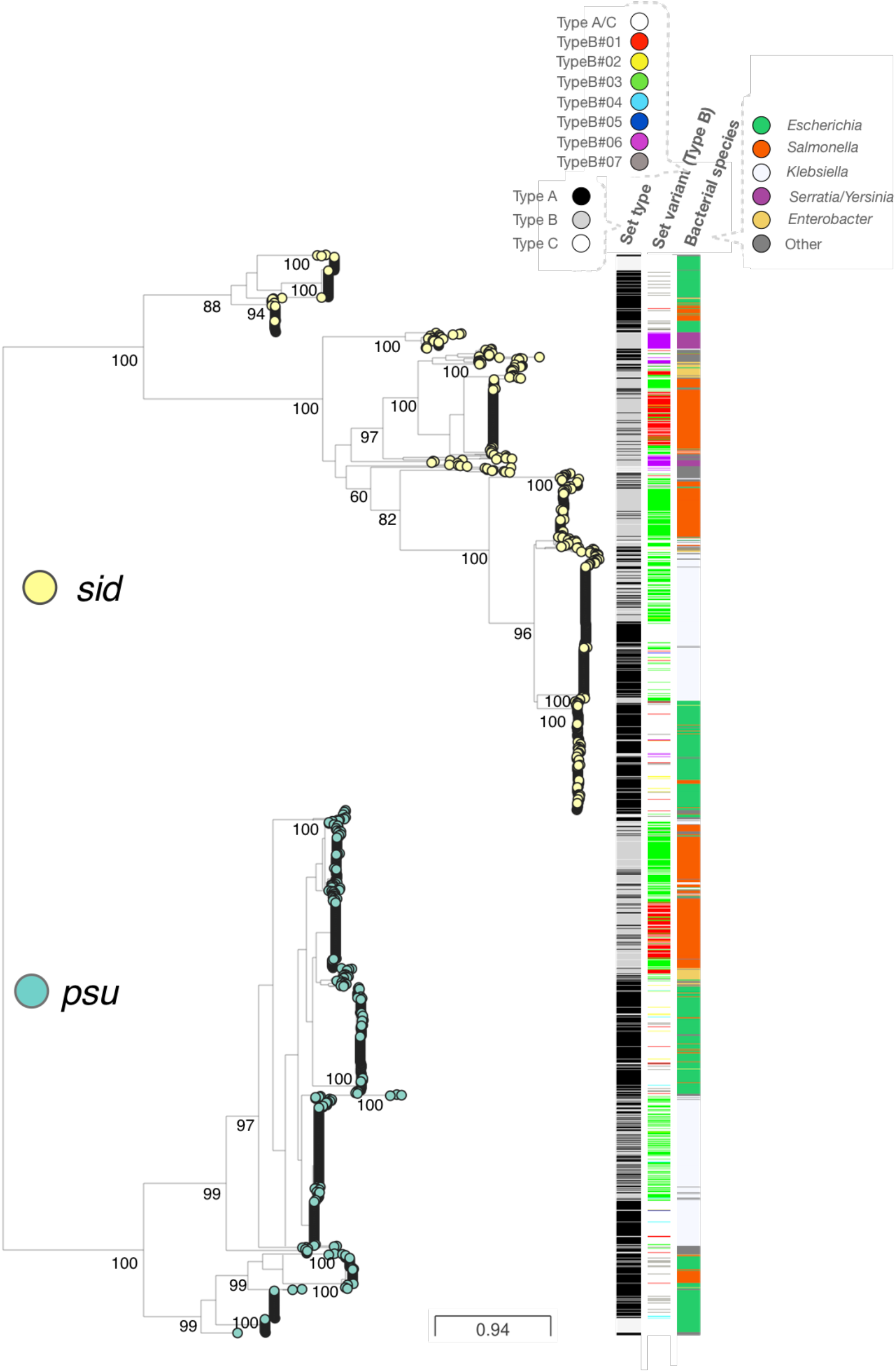
Joint phylogenetic tree of Psu and Sid proteins. The values shown on the branches are the result of ultrafast bootstraps and show a good support for the most important nodes. The columns on the right indicate, from left to right, the Type of set (A, B or C), the variant (only shown for sets of Type B, with Type A and all Type C variants shown as white spaces) and the bacterial species. The Newick tree file is included as File S3.

The sub-trees of Psu and Sid in the large joint phylogeny show a small early branching clade from elements present in *E. coli* and *S. enterica* that are those on the lower right cluster in the wGRR matrix (Fig 3). The Sid sub-tree then reveals a subsequent clade present in *Serratia*/*Yersinia* that is absent from the Psu tree. Hence, the ancestor of all P4-like satellites already contained the two key genes involved in the parasitism and elements of this clade lost *psu*.

The P4-like genomes of Types A or B are interspersed in the tree, often forming small separate clusters. This is consistent with the observations in the wGRR matrix and suggests that loss of one gene, typically *alpA*, still results in functional elements. The *psu* and *sid* homologs from Type C sets are either dispersed in the tree or clustered at the earlier branching clades of the sub-trees. The former may be genetically decaying P4-like satellites, but the latter might be part of different elements that have long diverged from P4 and lost *ε* and *α*. The phylogenetic tree is also consistent with the analysis of the wGRR matrix in that large clades of closely related proteins include a majority of elements from one bacterial genus and a few elements from other genuses. This reinforces the conclusion that these elements can transfer across large taxonomic groups of bacteria.

## Discussion

Recent experimental work has revealed the existence of phage satellites in Proteobacteria and Firmicutes [1]. Satellites of eukaryotic viruses are also known [27]. Such elements are thus likely to exist in other bacterial clades. There was some previous evidence of genetic diversity in PICI, for which more than twenty elements were described across diverse bacterial taxa [4]. Similarly, five different PLE were identified in *Vibrio cholerae* [10]. Yet, there was very little quantitative information on the frequency, gene repertoire diversity and organization, and on the taxonomic distribution of phage satellites in Bacteria. Here, we identified more than one thousand P4-like satellites across several genuses of Enterobacterales. They constitute a large family of characteristic mobile elements, with distinctive gene repertoires and a highly conserved genetic organization. The most sequenced bacterial species contain numerous such elements, e.g., they are found in almost half of the *E. coli* genomes, sometimes in multiple separate occurrences. These elements show very little homology with phages and are never found as plasmids. Hence, P4-like satellites are neither phages nor plasmids, but instead a distinct and widespread lineage of integrative mobile genetic elements.

Some P4-like genomes miss one or, in rarer cases, two core genes raising the possibility that they are defective. Many prophages are non-functional in bacteria [28,29] and it is possible that some of the P4-like satellites are also non-functional. Yet, three pieces of evidence suggest that most Type B and C sets are functional variants of P4. First, some genes are almost always present whereas one, *alpA*, is more often absent. This gene might be a gene expression regulator [4], which are often replaced in mobile elements. If Type B and C elements were non-functional, one would expect a more random distribution of absent genes. We found that one gene present in most P4-like elements is actually lacking in P4, in agreement with the idea that some of these genes can be missing in functional elements. Second, for at least two Type B variants (those lacking *psu* or *alpA*), the position of the missing component rarely contained a pseudogene of the missing gene (8% of *psu*-less variants, and less than 0.1% of *alpA*-less variants), suggesting that the majority of these elements are not recent loss-of-function variants. Third, many of the Type B and C elements form small conserved clades within the Psu+Sid phylogenetic tree, suggesting that they are functional. One particularly intriguing case concerns the elements that have lost *psu*, including those among *Yearsineacea*. Since *psu* and *sid* are homologs, *psu*-less elements could rely on *sid* variants that are able to single-handedly sequester the capsid of helper phages. Overall, while we cannot ascertain the function of all elements that we have identified, many seem to contain a full or nearly full repertoire of the key components of the family and may therefore be functional.

The origins of P4-like elements are unknown. We observed very little sequence similarity between phages and P4-like elements, suggesting they had distinct evolutionary origins. Most of these homologs concern genes that are frequently transferred across mobile genetic elements like replicases, integrases and transcriptional regulators. Accordingly, the integrase phylogenetic tree is very different from the others. This suggests frequent gene exchanges with other mobile genetic elements and is consistent with the observation above that the elements tend to be scattered across the genome. This is also consistent with the proposal that the *cos* site, between the integrase and *psu*, is associated with rapid diversification of the defense systems carried by P4-like elements [9]. *δ* is the only specific P4 gene that has clear homologs in the *ogr* genes of P2-like phages, but sequence similarity is very low [3]. P2-like phages are probably also very ancient [30] and may have co-evolved with P4-like elements for a long time. The evolutionary history of the two most distinctive P4 genes – *psu* and *sid* – should be key to trace the origins of P4. However, these genes do not show significant similarity with phage genes and while Psu and Sid have similar protein structures, they are very dissimilar from all other known protein structures [18,31]. It is tempting to speculate that *psu* and *sid* originated by gene duplication (or gene transfer leading to an equivalent outcome) followed by specialization into a protein that forms a transient external scaffolding cage around the P4 procapsids (Sid) and another acting as a stabilizing protein at the outside of P4 capsids (Psu). Given the genetic diversity of the components of P4-like elements, their variation in gene repertoires, their broad distribution across Enterobacterales and the differences between Psu and Sid, P4-like elements are probably a very ancient family of phage satellites.

The impact of P4-like satellites depends on their ability to disseminate, the traits they carry and how they affect the helper phages. The high incidence of very similar P4-like satellites in different bacterial genuses suggests they have a broad host range. If the host range of the satellite depends on the ability of the virion to attach and inject DNA in a recipient cell, then it should be limited by the host range of its helper phage. However, satellites can increase their host range if they can exploit multiple helper phages. This may be facilitated by the acquisition of genes at the elements’ hotspots, which have very diverse gene repertoires contributing for the open pangenome of P4-like elements. These regions also include traits that affect bacterial fitness, such as the anti-phage defense systems identified in a P4 hotspot [9]. In order to better understand the basis for the variation and distribution of P4-like satellites in bacterial species, we will now analyze their co-incidence with P2-like prophages. This will pave the way to study the population and co-evolutionary dynamics of satellites and their helper phages. Our novel collection of P4-like elements will thus facilitate the study of the impact of phage satellites in the diversification of bacterial genomes and in phage-bacteria dynamics.

## Materials and methods

### Data

We retrieved the complete genomes of 13513 bacteria, 22310 plasmids and 2502 phages from NCBI non-redundant RefSeq database (ftp://ftp.ncbi.nlm.nih.gov/genomes/refseq/, last accessed in May 2019). Five phage genomes were excluded from the analysis because of lack of gene annotation, resulting in a dataset with 2487 phage genomes. Prophages (integrated temperate phages) were predicted in bacterial genomes using VirSorter v.1.0.3 with the RefSeqABVir database [32]. The least confident predictions, i.e., categories 3 and 6, which may be prophage remnants or erroneous assignments, were excluded from the analyses, resulting in a total of 26984 prophages.

### Detection of P4-like elements using reference components

We used HMMER (v3.1b2 [33]) to search for homologs of each component (PFAM or custom profiles in parenthesis): Psu (PF07455.12), δ (PF04606.13), Sid (custom profile, in Supplementary File S2), AlpA (PF05930.10), ε (PF10554.10), α (PF03288.17) and integrases (PF00589.20, for Tyrosine recombinases, and PF00239.19 and PF07508.11 together for Serine recombinases) in bacterial and plasmid replicons. We retained the hits with an e-value of at most 10^-5^ and a profile coverage of at least 40%. We then ordered the genes by their position in each replicon and searched for consecutive pairs of components less than 10 open reading frames apart (see Fig S2). Additional components were added by transitivity when they were within 10 ORFs of the set. This was repeated until either the following component was at a distance larger than 10 ORFs (which would start a new process of constructing a different set, closing the current one), or when there were no more components in the replicon. The sets sometimes contain duplicates of a given component, particularly integrases which tend to aggregate in bacterial genomes at chromosome hotspots [34]. When more than one integrase was part of the set, we kept the integrase closest to *psu*, as expected from the linearization of the original P4 genome, and our analyses confirm the conservation of this location for the integrase. When components were in multiple copies, which is very rare, we kept the first occurrence.

### Pangenomes of the P4-like satellite family

P4-like satellites of Types A and B were delimited by the integrase and the furthest component of the set (in the case of the variant TypeB#07, which lacks the integrase, we used *psu* since it is the component most frequently found after the integrase). We computed the pangenome of the P4-like elements by clustering all their proteins at a minimum of 40% identity, using mmseqs2 [35] (Nature Biotechnology release, August 2017), with parameters –cluster_mode 1 and –min_seq_id 0.4 (all other parameters were left as default). The resulting gene families were functionally annotated using PFAM (release 33.1).

### Weighted gene repertoire relatedness between P4-like genomes

We searched for sequence similarity between all proteins of all phages using mmseqs2 (Nature Biotechnology release, August 2017 [35]) with the sensitivity parameter set at 7.5. The results were converted to the blast format and we kept for analysis the hits respecting the following thresholds: e-value lower than 0.0001, at least 35% identity, and a coverage of at least 50% of the proteins. The hits were used to retrieve the bi-directional best hits between pair of phages, which were used to compute a score of gene repertoire relatedness weighted by sequence identity:

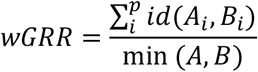

where A_i_ and B_i_ is the pair *i* of homologous proteins present in *A* and *B*, id(*A_i_*,*B_i_*) is the sequence identity of their alignment, and min(*A*,*B*) is the number of proteins of the genome encoding the fewest proteins (*A* or *B*). wGRR is the fraction of bi-directional best hits between two genomes weighted by the sequence identity of the homologs. It varies between zero (no bi-directional best hits) and one (all genes of the smallest genome have an identical homolog in the largest genome). wGRR integrates information on the frequency of homologs and sequence identity. For example, when the smallest genome has 10 proteins, a wGRR of 0.2 can result from two homologs that are strictly identical or five that have 40% identity. The hierarchical clustering of the wGRR matrix, and the corresponding heatmap, were computed with the *clustermap* function from the *seaborn* package (version 0.9), using the *average* (UPGMA) clustering algorithm.

### Join phylogenetic tree of *psu* and *sid* components

We concatenated all protein sequences of *psu* and *sid* from all sets of Types A, B and C, including also the respective components from the original P4 genome (NC_001609). The sequences were aligned using mafft-linsi [36] (version 7.222, default parameters) and the resulting alignment trimmed with noisy [37] (version 1.5.12, default parameters). We used IQ-TREE [38] (version 1.6.5) to build the phylogenetic trees, with the options –wbtl (to conserve all optimal trees and their branch lengths), –bb 1000 to run the ultrafast bootstrap option with 1000 replicates,−m MFP for automatic model selection, and –nt AUTO. The resulting tree files were converted to Newick format using the Phylogeny.fr [39] webserver (http://phylogeny.lirmm.fr/phylo_cgi/index.cgi, last accessed March 2021). The visualization of the tree was performed using the Microreact [40] webserver (https://microreact.org/, last accessed March 2021).

### Analysis of the core genome and tree of *E. coli*

The pangenome, core genome and the tree of *E. coli* were computed with Panacota [23]. Briefly, the genomes of *E. coli* in our dataset were retrieved, filtered to remove very closely related genomes (MASH distance < 0.0001), and then clustered using mmseqs2 with a minimal threshold of 80% identity in protein sequences. Within the pangenome of these 657 genomes there were 2107 families present in more than 90% of the genomes, which make the core genome. The core genome was aligned (for families with not more than a single gene per genome) using mafft-linsi, rendering a multiple alignment with 2069583 positions that was used to build a phylogenetic tree by FastTree 2.1 with model JC [24]. The position of P4-like elements in *E. coli* was computed in relation to the core genome. The genes of the core genome were positioned as in the most ancient genome of *E. coli* (strain MG1655). For each P4-like element we identified the immediately upstream gene family of the core genome in the focal genome. If the core gene family has an ordered position in MG1655 of N, we noted N+1 for the position of the P4-like element. We then plotted the histogram of the positions.

## Acknowledgements

We thank José Penadés and Marie Touchon for comments and suggestions on earlier versions of the manuscript. We acknowledge the financial support of Equipe FRM (EQU201903007835), Laboratoire d’Excellence IBEID (ANR-10-LABX-62-IBEID) and the ANR (SALMOPROPHAGE ANR-16-CE16-0029).

## Data availability

The bacterial and phage genomes, as well as most profiles used to detect the core components of P4, are publicly available. The custom profile corresponding to sid is included as supplementary file. The code used for the analysis will be made available upon request to the authors.

