## Supplementary Data for "To catch a hijacker: abundance, evolution and genetic diversity of P4-like bacteriophage satellites"

**This PDF file includes:**

Figures S1 to S9

Legends for Files S1 to S7

**Other supplementary materials for this manuscript include the following:**

Files S1 to S7

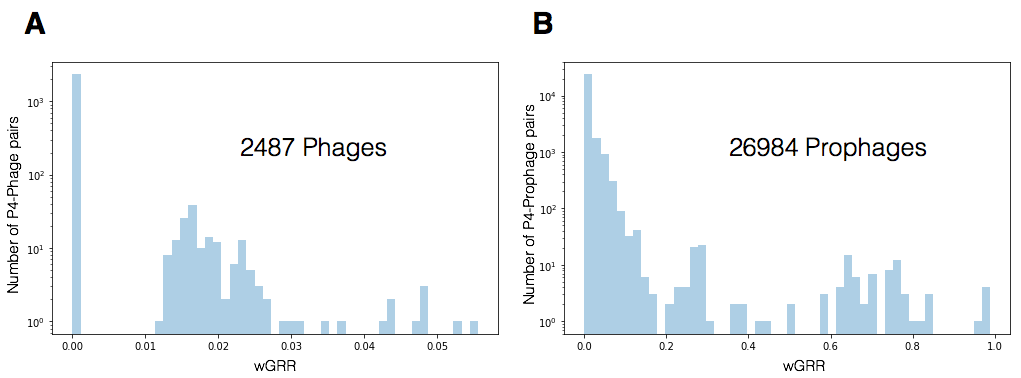

**Figure S1. Genetic similarity of P4 with phage and prophage databases.** Shown are the distributions of the values of wGRR between P4 and elements from **A)** the dataset of phage sequences in NCBI, and **B)** the prophages in bacterial genomes identified with Virsorter. Note the log scale in the Y axis.

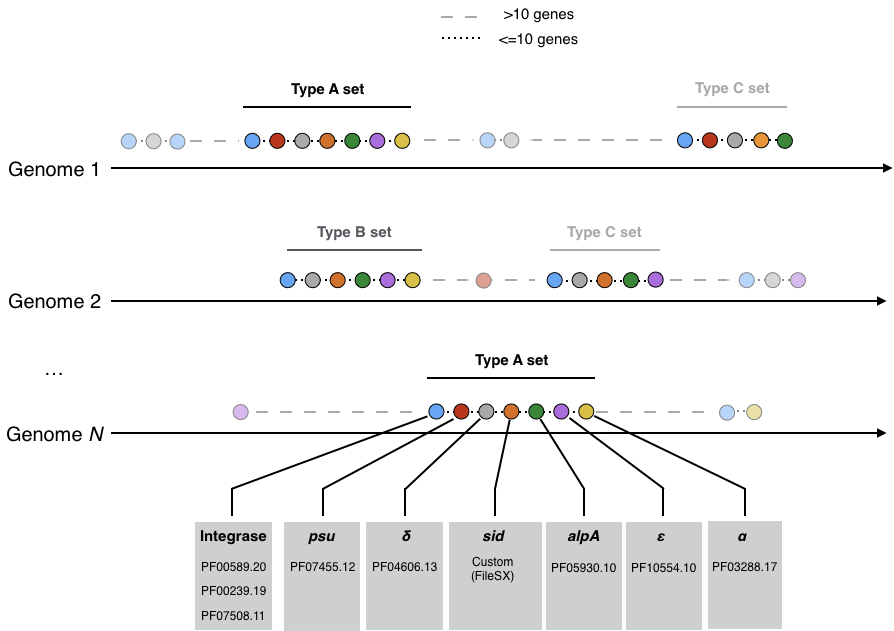

**Figure S2. Detection of complete and incomplete sets of P4 core genes in bacterial genomes.** Homologs of core genes were detected in bacterial and plasmid genomes using the HMM profiles indicated in grey boxes. Sets were constructed from consecutive pairs of homologs less than 10 genes apart. If the closest homolog is at a larger distance, the current set is closed and a new one starts. Sets of Type A have homologs of all the seven core genes. Sets of Type B and C lack one or two of the components, respectively.

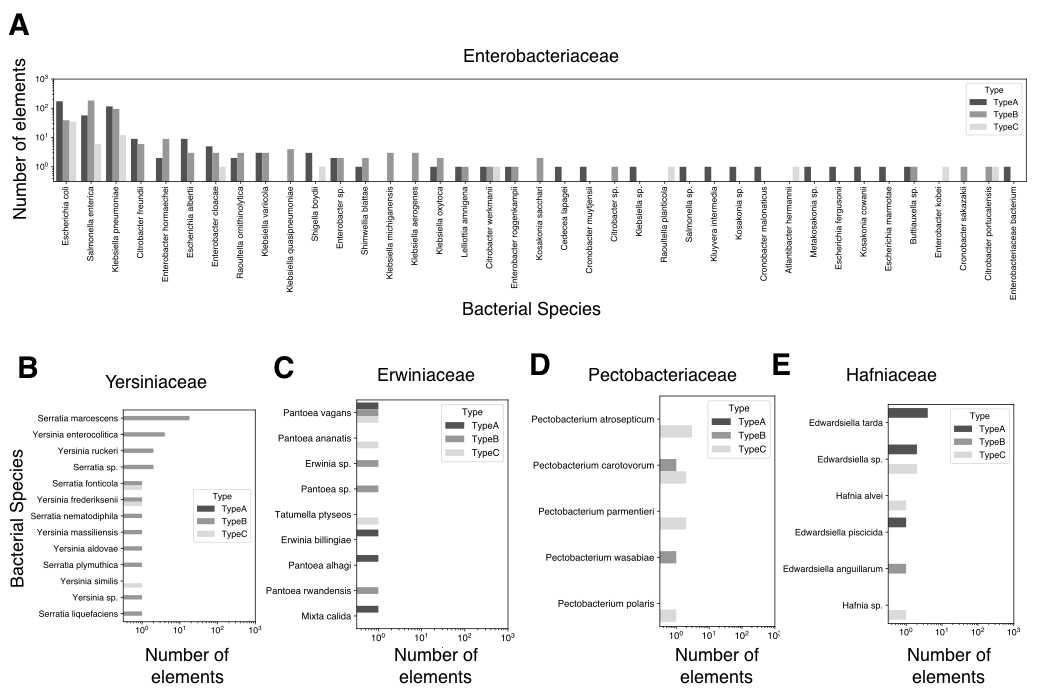

**Figure S3. Number of P4-like elements in the bacterial species from different families.** Differentially shaded bars indicate sets of Type A, B or C (from darker to lighter shades).

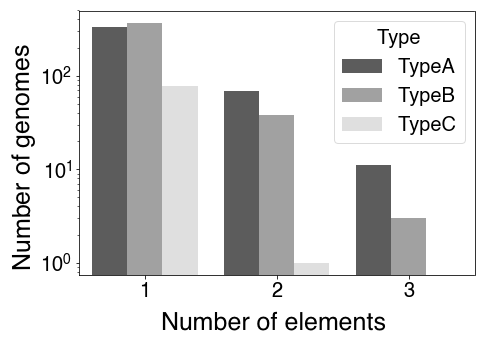

**Figure S4. Number of P4-like elements per bacterial genome.** Differentially shaded bars indicate sets of Type A, B or C (from darker to lighter shades).

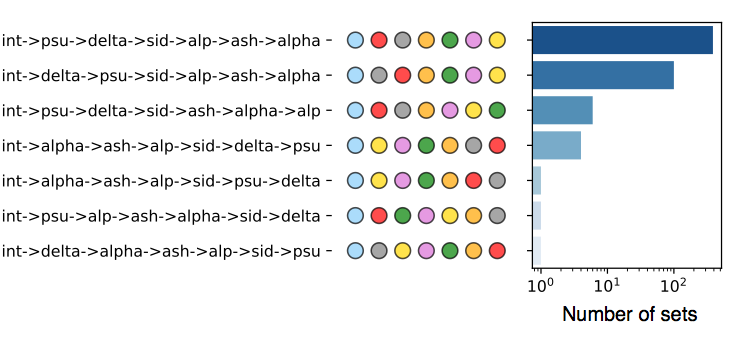

**Figure S5. Organizational variants of P4-like genomes of Type A.** The circles on the left indicate the organizational variant, and the bars on the right indicate (in log scale) their frequency amongst Type A P4-like genomes.

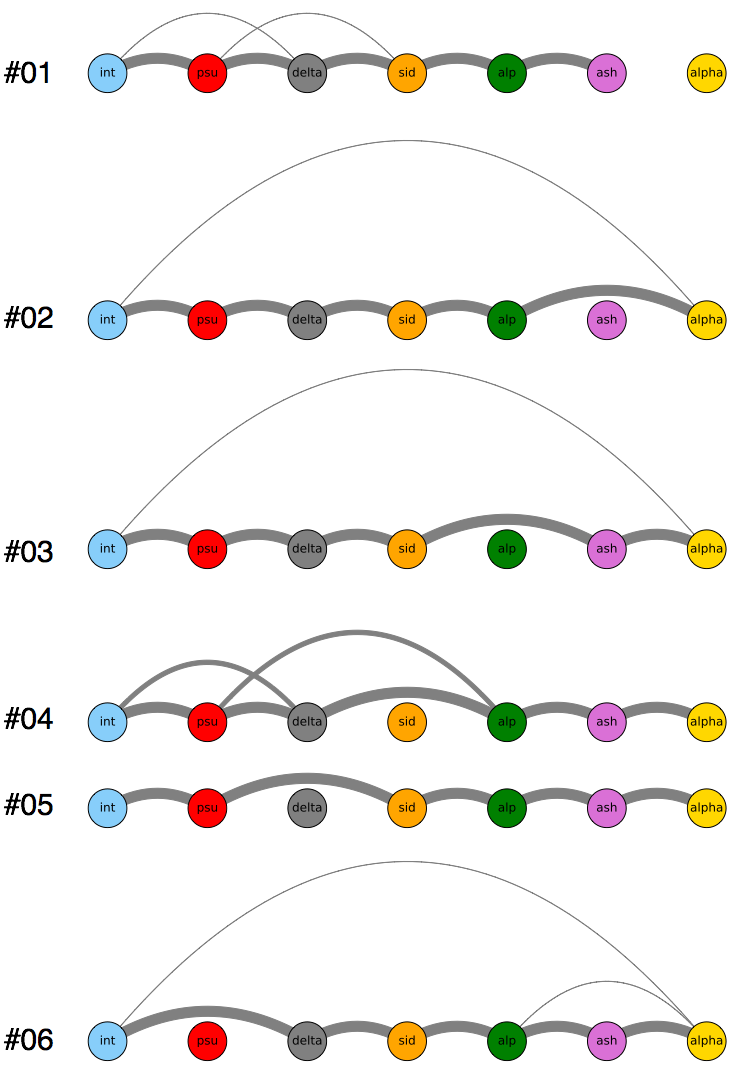

**Figure S6. Genetic organization of P4-like sets of Type B.** Shown are the organization of core genes found in the different variants (excluding the one lacking the integrase) of Type B sets. The thickness of the edges is proportional to the frequency of adjacency of pairs of components. The component missing in each variant was kept visible to maintain consistency across the diagrams, but are not linked to any other component(s).

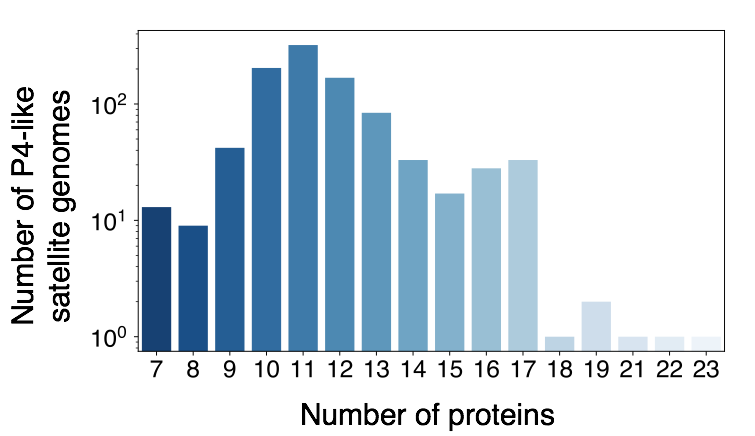

**Figure S7. Distribution of the number of proteins in P4-like genomes.**

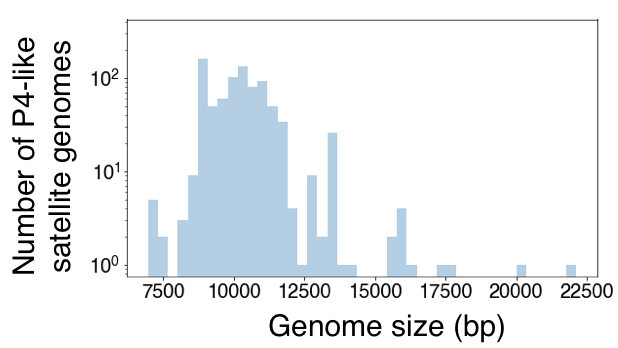

**Figure S8. Genome size distribution of P4-like genomes, excluding Type B variants lacking the Integrase or *α*, which are usually at the edges of the elements in the bacterial genome.**

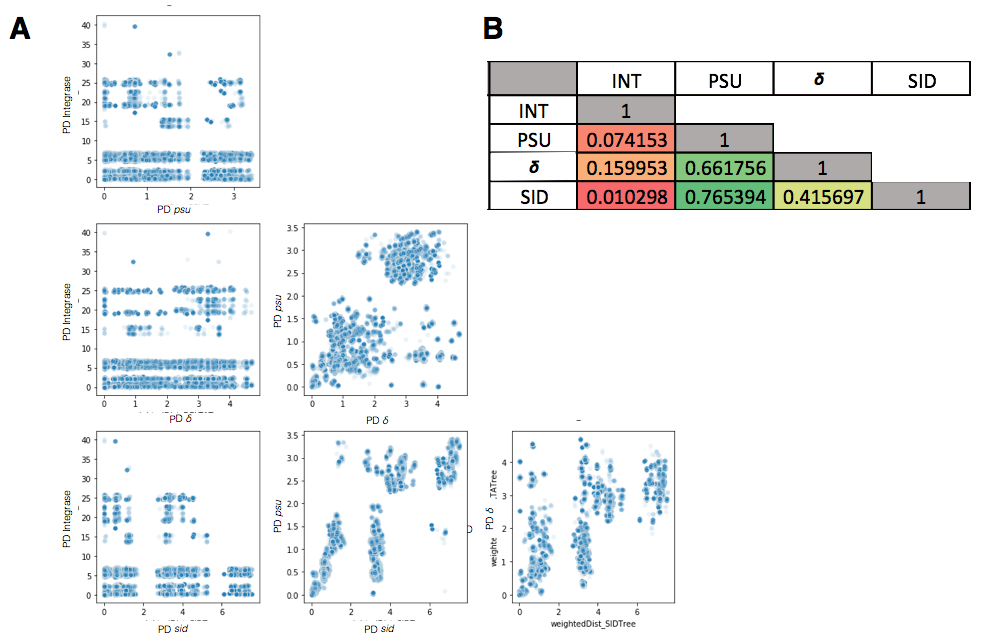

**Figure S9. Phylogenetic similarity between the Integrase, Psu, *δ* and Sid of P4-like genomes.** Phylogenetic similarity was computed by comparing the Patristic Distances (as a sum of branch lengths) between all the leaves that are common between the phylogenetic trees of the Integrase, Psu, delta and Sid (897 P4-like genomes in total). **A)** Scatterplots of the Patristic Distances for the same pairs of leaves between the different components, indicated in the axis of each panel. **B)** Spearman correlation values for the scatterplots shown in **A**. Color code shows the strongest (greenest) to weakest (reddest) correlation values.

File S1 (separate file). Table with all the 1037 P4-like sets (Types A, B and C) found in bacterial genomes.

File S2 (separate file). Hidden Markov Models for *sid*.

File S3 (separate file). Phylogenetic tree (Newick format) of Psu and Sid

File S4 (separate file). Phylogenetic tree (Newick format) of Integrase

File S5 (separate file). Phylogenetic tree (Newick format) of Psu

File S6 (separate file). Phylogenetic tree (Newick format) of 𝛿

File S7 (separate file). Phylogenetic tree (Newick format) of Sid
